# Somatic Mutations and Neoepitope Homology in Melanomas Treated with CTLA-4 Blockade

**DOI:** 10.1101/088286

**Authors:** Tavi Nathanson, Arun Ahuja, Alexander Rubinsteyn, Bulent Arman Aksoy, Matthew D. Hellmann, Diana Miao, Eliezer Van Allen, Taha Merghoub, Jedd Wolchok, Alexandra Snyder, Jeff Hammerbacher

## Abstract

Immune checkpoint inhibitors are promising treatments for patients with a variety of malignancies. Toward understanding the determinants of response to immune checkpoint inhibitors, it was previously demonstrated that somatic mutation burden is associated with benefit and a hypothesis was posited that neoantigen homology to pathogens may in part explain the link between somatic mutations and response. To further examine this hypothesis, we reanalyzed cancer exome data obtained from a previously published study of 64 melanoma patients treated with CTLA-4 blockade and a new dataset of RNA-Seq data from 24 of those patients. We found that the predictive accuracy does not increase as analysis narrows from somatic mutation burden to predicted MHC Class I neoantigens, expressed neoantigens, or homology to pathogens. Further, the association between somatic mutation burden and response is only found when examining samples obtained prior to treatment. Neoantigen and expressed neoantigen burden are also associated with response, but neither is more predictive than somatic mutation burden. Neither the previously-described tetrapeptide signature nor an updated method to evaluate neoepitope homology to pathogens were more predictive than mutation burden.

## Introduction

Checkpoint blockade therapies are improving outcomes for patients with metastatic solid tumors (1–4). As only a subset of patients respond, there is a critical need to identify determinants of response. Expression of program death-ligand one (PD-L1) is the lead companion diagnostic for PD-1/PD-L1 blockade therapies, but sensitivity and specificity are limited (5–7). Recent studies have demonstrated an association between elevated tumor mutation burden and benefit from checkpoint blockade therapies (8–11).

In a recent study of melanomas treated with checkpoint blockade agents targeting cytotoxic T-lymphocyte associated protein 4 (CTLA-4) (8) the authors present the hypothesis that responding tumors may share features with each other or with infectious agents, and that such resemblance may predict response. In the present report, we performed a reanalysis of the data in that study using updated methods and integrating new RNA sequencing data from a subset of 24 samples.

We found that in this small dataset, nonsynonymous mutation burden was associated with clinical benefit from therapy in samples collected prior to, but not after treatment with CTLA-4 blockade. Predicted neoantigen burden and percentage of C to T transitions characteristic of ultraviolet damage were associated with, but did not outperform mutation burden. We developed a publicly available tool, Topeology (https://github.com/hammerlab/topeology) to compare neoantigens to known pathogens. Neither the resemblance of tumor neoantigens to known antigens, nor the previously published tetrapeptide signature outperformed mutation burden as a predictor of response.

## Materials and Methods

### Patient Samples

All analyzed samples were collected in accordance with local Internal Review Board policies as described in (8) and summarized in Table 1. 34 patients had tumor samples collected prior to initiating CTLA-4 blockade and 30 patients had samples collected after initiating CTLA-4 blockade. Clinical benefit was defined as progression-free survival lasting for greater than 24 weeks after initiation of therapy. 9 discordant lesions were present, where overall patient benefit did not match individual tumor progression. See Table 1 for a cohort summary and Supplementary Information (“Clinical patient data”) details about this patient cohort.

**Table 1.** Cohort Summary. Features of tumors from patients with clinical benefit, no benefit, or in which a discordant lesion was resected.

| Group | Benefit | No Benefit | Discordant |
| --- | --- | --- | --- |
| **N** | 27 | 28 | 9 |
| **% Cutaneous** | 20/27 | 19/28 | 5/9 |
| **OS** | 3.7 (1.6 - 7.3) | 0.8 (.2 - 2.7) | 4 (1.7 - 7.9) |
| **Age** | 65 (33 - 81) | 58.5 (18 - 79) | 68 (40 - 90) |
| **Mutations** | 611 (165 - 3394) | 321 (6 - 1816) | 549 (93 - 1336) |
| **Neoantigens** | 1388 (209 - 6502) | 714.5 (3 - 4510) | 1048 (197 - 2584) |

### Mutation Calls

Single nucleotide variants (SNV) were called with an ensemble of four variant callers: Mutect, Strelka, SomaticSniper, and Varscan as previously described (9). Insertions and deletions (indels) were called using Strelka with default settings.

### HLA Typing

HLA types were determined by *ATHLATES* for all samples using exome sequence data and confirmed with *seq2HLA* for samples that had RNA-Seq available (24 samples).

### Neoepitope Prediction

Somatic SNVs that occurred a single base away from other somatic SNVs were combined into larger variants containing both SNVs. For each somatic variant we used Topiary (<u>https://github.com/hammerlab/topiary</u>) to generate the predicted 8-11mer amino acid product resulting from somatic alterations (SNV or indel), including predicted neoepitopes generated from combined SNVs. Each variant was linked to its corresponding coding DNA sequence (CDS) from Ensembl based on its B37 coordinates. The CDS sequence was re-translated with the mutated DNA residue producing the mutated peptide product. NetMHCcons v1.1 generated a predicted binding affinity for all 8-11mers containing the mutated amino acid and all peptides with an IC50 score below 500nM were considered predicted neoepitopes. For variants longer than a single residue we looked at all 8-11mers generated downstream of the variant. Neoepitopes from (10) were generated from a separate pipeline as published.

### RNA-Seq

The 24 tumor RNA samples were a subset of the published 64-sample dataset, and included those samples from the 64-sample set that had sufficient tissue for RNA isolation. Data from some of samples have been previously presented (12). Sequencing libraries were prepared from total RNA with the Illumina TruSeq mRNA library kit (v2) and then sequenced on the HiSeq 2500 with 2x50bp paired reads, yielding 47-60M reads per sample (New York Genome Center). The RNA reads were aligned using the STAR aligner after which Cufflinks was used for gene quantification (FPKM). seq2HLA was also used to quantify HLA gene expression (RPKM). Allele-specific expression was measured by the fraction of RNA reads supporting the variants found in exome sequencing.

Gene set enrichment analysis (GSEA) was performed using v2.2.0 of the software provided by Broad Institute (<u>http://www.broadinstitute.org/gsea/index.jsp</u>) and the Hallmark gene set collection that was used in the comparison was accessed on August 18, 2015 from the MSigDB website (<u>http://www.broadinstitute.org/gsea/msigdb/index.jsp</u>). The Hallmark gene set collection was extended by adding gene symbols corresponding to well-known peptides that are (i) tumor-specific; (ii) associated with differentiation; (iii) overexpressed in cancer cells. To do this, gene symbols were imported from the Cancer Immunity Peptide Database as gene sets that are compatible with the GSEA software. Before running the GSEA, the gene expression data (FPKM) were collapsed using official gene symbol identifiers and the median expression value used when multiple transcripts mapped to the same gene symbol. To normalize the data further, non-informative genes with no variation (standard deviation of 0) across all samples were removed. Three GSEA analyses were conducted comparing: (i) pre-treatment benefiting vs. pre-treatment non-benefiting; (ii) pre-treatment benefiting vs. post-treatment benefiting; (iii) pre-treatment non-benefiting vs. post-treatment non-benefiting. In all these comparisons, the normalized gene expression values were used as the input matrix. The number of permutations was set to 1,000, restricting these permutations to gene set labels rather than the sample phenotype labels due to our sample size, and we kept the rest of the default options (see <u>http://www.hammerlab.org/melanoma-reanalysis/gsea-results/</u> for complete reports and instructions to replicate them).

### Neoantigen Homology

We developed a tool, Topeology, to compare tumor neoepitopes to entries in the Immune Epitope Database (IEDB) (13), accounting for position, amino acid gaps, and biochemical similarity between amino acids. Epitopes were compared, and the comparison scored, using the Smith-Waterman alignment algorithm (14) supplied with a substitution matrix consisting of PMBEC correlation values derived from the PMBEC covariance matrix (15). We compared amino acids from position 3 to the penultimate amino acid of the peptide, assuming that the anchor residues would be necessary for MHC Class I presentation and would therefore not be “visible” to a T cell. A gap penalty equal to the lowest PMBEC correlation value was supplied (Figure 3B). Peptide comparisons were only considered if they were the same length. Smith-Waterman scores were normalized for length by dividing by the length of the peptide section being compared (Figure 3B).

Peptides were only included for comparison if the mutant peptide score was greater than or equal to the wild type peptide score. In addition, an epitope from IEDB was only compared to a neoepitope if either (i) the patient’s HLA allele(s) presenting that neoepitope were listed as HLA alleles for the IEDB epitope or (ii) the IEDB epitope was a predicted binder for one of the patient’s alleles.

In order to narrow down candidate epitopes to those with some evidence of biological relevance, peptides from IEDB were required to exhibit *in vitro* evidence of human T cell activation. Because many peptides showed different T cell responses depending on the assay used, at least 60% of the instances of that specific amino acid sequence found in IEDB were required to exhibit an activated T cell response in order for that peptide to be considered “T cell activating.” Peptides were required to be 8 to 11 amino acids in length and were filtered to remove allergens, zoonotic organisms not known to affect humans, and self-epitopes. Limited manual filtering of source organisms was also performed to ensure the exclusion of non-pathogenic antigens. Peptides from IEDB were evaluated both as a whole and as two groups: viral and nonviral pathogens.

### Predictive Model Evaluation

To evaluate whether similarity to known pathogenic antigens can predict clinical benefit, we generated predictive models using logistic regression with 𝓁1 regularization. Only pre-treatment samples were considered, and a single pre-treatment sample that was considered discordant was also excluded. One model attempted to predict clinical benefit for each sample based on the maximal similarity of the sample’s neoepitopes to each IEDB epitope, with a feature for every IEDB epitope. Another model did the same, but aggregated IEDB epitopes based on their source pathogens, resulting in a feature for every IEDB source pathogen. The maximal similarity to an epitope (or pathogen) is referred to as the “score” for that epitope (or pathogen) in Results. To generate two additional models, IEDB epitope features were averaged together to create a single-feature model, and the same method was applied to IEDB pathogen features. See Supplementary Information for a table of all models, including feature counts per model.

We tested these predictive models using 1,000 rounds of bootstrapping in order to generate stable measures of performance, using 75% of the samples for training. For each round, we performed 100 inner rounds of bootstrapping to optimize the regularization strength and scaling hyperparameters using 75% of the outer training samples for hyperparameter fitting and 25% of the outer training samples for hyperparameter validation. Each inner and outer round of bootstrapping calculated the area under the curve (AUC) of the receiver operating characteristic. A baseline AUC, which was also calculated using the same bootstrapping procedure, used mutation burden in place of a logistic regression probability. Each of the 1,000 AUC values generated by the outer bootstrap sampling was compared with the corresponding baseline mutation burden AUC for the same sampling, resulting in a distribution of differences between each pair. Confidence intervals, as described by (16) were taken from these AUC distributions and pairwise AUC difference distributions.

We also created the same similarity scores and predictive models for the neoepitope predictions generated by (10). Because the pipelines were different, as well as the definitions of clinical benefit, these results are not fully comparable and are found in the supplementary materials (Supplementary Information).

### Tetrapeptide Signature Evaluation

We evaluated the tetrapeptide signature approach from Snyder et al (8) using the above bootstrapping procedure. Because Snyder et al used validation data to impact signature creation, we did not validate the identical tetrapeptide signature generated in (8). Instead, in order to perform validation, we generated additional tetrapeptide signatures from discovery data alone, excluding any validation set filtering. We used the same candidate tetrepeptide generation approach used in (8) as opposed to our updated approach used above: positionality was not considered, HLA alleles were not considered, and IEDB peptides were not filtered by length. We repeated the analysis twice: once using the variant calls and cohort (n=64) from (8) and once using updated variant calls and only pre-treatment, non-discordant samples (n=33).

The bootstrapping procedure considered 1,000 randomly sampled training sets. The signature rules from Snyder et al, which are as follows, were applied to each sampled discovery set to generate separate signatures: a tetrapeptide was added to the signature if it was present in either (i) neoepitopes from at least 3 different benefit patients or (ii) neoepitopes from 2 benefit patients and a T cell activating epitope from IEDB. Tetrapeptides that occurred in non-benefit patient neoepitopes were excluded from the signature. For each sampling round, these signature rules only considered the round’s discovery set so that each of the 1,000 signatures generated could be tested against their associated validation sets.

Performance was measured using a single binary value, as described by Snyder et al: whether or not a patient’s neoepitopes contained any of the signature tetrapeptides. In this binary case, the receiver operating characteristic curve contains a single threshold. The AUC score defined by the area under the line segments that connect to this single threshold is equal to balanced accuracy, which is the metric we used in this case. We also evaluated the AUC using per-patient counts of signature tetrapeptides.

## Results

### Mutation and Expressed Mutation Burden are Associated with Outcome in Pre-Treatment Samples in Advanced Melanoma Patients Treated with CTLA-4 Blockade Using Updated Bioinformatic Analysis

We reanalyzed the mutation burden of the melanoma tumors included in Snyder et al (8) using a modified system of four callers (as described in (9)). Analyzing the data using this system increased the median nonsynonymous mutation burden of the group 1.9-fold from 248 to 471 (original range 1-1878, to new range 6-3394) (Supplementary Figure 1A-B).

In samples collected prior to treatment (n=34), mutation burden was higher in patients with clinical benefit (Figure 1A, median: 654 in benefiters versus 196.5 in non-benefiters, Mann-Whitney, p=0.0006), and elevated mutation burden was associated with overall survival (Supplementary Figure 1C, log-rank test, p=0.01). Of pre-treatment samples, one patient who otherwise experienced disease control had a progressing lesion resected, representing a discordant lesion (Methods). In patients whose tumor samples were collected after initiating CTLA-4 blockade (n=30), there was not a significant difference in the mutation burden between benefit and non-benefit groups (Figure 1B, median: 592 in benefiters versus 396 in non-benefiters, Mann-Whitney, p=0.19), and elevated mutation burden was not significantly associated with overall survival (Supplementary Figure 1D, log-rank test, p=0.29). 8 discordant lesions were present among post-treatment samples; when excluding patients with discordant lesions, there was still not a significant difference in the mutation burden between benefit and non-benefit groups (median: 592 in benefiters versus 392 in non-benefiters, Mann-Whitney, p=0.20) or a significant association with overall survival (log-rank test, p=0.39). In sum, mutation burden was associated with clinical benefit only in samples collected prior to treatment.

**Figure 1.**
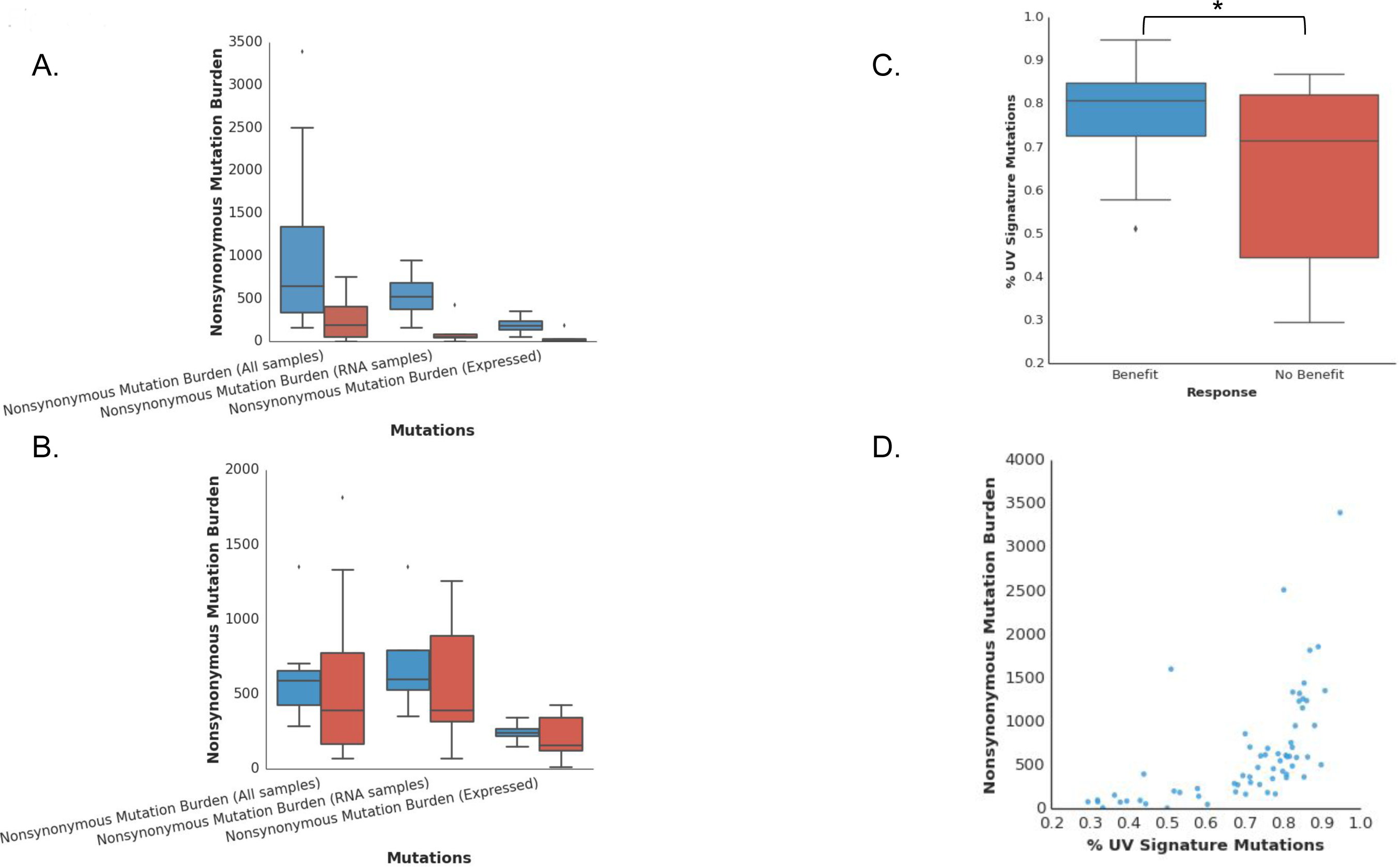
Mutation Burden and Ultraviolet Signature. A. Median and range of mutation burden and allele-specific expression of mutations in samples collected prior to treatment (for benefit versus no benefit, all (n=34, Mann-Whitney, p=0.0006), expressed (n=9, p=0.024)). In A and B, the first pair of bars represent mutation burden for all sequenced tumors; the second pair represent mutation burden in the subset of tumors for which RNA was available; the third represent the expressed mutations. In A-C, blue bars represent benefiting tumors; red bars represent non-benefiting tumors. B. Median and range of mutation burden and expressed mutations in samples collected after treatment (for benefit versus no benefit, all (n=30, Mann-Whitney, p=0.19), expressed (n=15, p=0.46)). C. Correlation between signature of DNA damage from ultraviolet (UV) exposure and clinical response (*Mann-Whitney, p=0.003). D. Correlation between UV signature and mutation count (Spearman rho=0.77, p=4e-14).

In some previously published studies, insertion or deletion mutations (indels) have not been considered in calculating mutation and neoantigen burdens. Genetic alterations of greater than one amino acid can theoretically generate peptides that are substantially different from the wild type as a result of a shifted reading frame (termed neo-open reading frames, or neo-ORFs (17)). The number of predicted neoantigens resulting from indels was also associated with outcome (median 9 and 6 in benefiters and non-benefiters, respectively, Mann-Whitney, p=0.018), but comprised a very small minority of neoantigens (median of 0.8% of all predicted neoantigens, range 0-33.7%). The possibility remains that these transcripts or the resulting translated proteins are subsequently degraded (18).

The previously published analysis did not find a correlation between the signature of ultraviolet DNA damage and clinical benefit. We reexamined this question using updated methods. A tumor was determined to have a UV signature when > 60% of mutations were C>T transitions at dipyrimidine sites (19,20). As expected, 36 of 44 (81%) tumors of cutaneous origin harbored the UV signature. Five out of six tumors with acral or uveal histology did not have a UV signature (although one did: ID 6819). In contrast to the originally published data, we found the rate of UV signature mutations correlated with clinical benefit (Figure 1C, Mann-Whitney, p=0.003) and overall mutation burden (Figure 1D, Spearman rho=0.77, p=4e-14).

Two studies have investigated expressed neoantigens in human samples from patients treated with immunotherapy by examining the expression level of genes which harbored mutations (10,21). However, because many mutations do not result in expressed proteins, we examined allele-specific expression (see Methods) of mutations. The median rate at which genes containing mutations were expressed (FPKM > 0) in all samples was 37% (range 20%-50%). Of all tumors with available RNA-Seq data (n=24), one post-treatment lesion was discordant. For tumors sampled prior to treatment with available RNA-Seq data (n=9), patients with long-term benefit had a higher number of mutations expressed in the RNA samples (Figure 1A, Mann-Whitney, p=0.002). For tumors sampled after treatment (n=15), the difference in expressed mutation burden between benefiters and non-benefiters was not significant (Figure 1B, Mann-Whitney, p=0.46). Predicting clinical benefit using expressed mutation burden among pre-treatment samples with RNA-Seq data (n=9) resulted in an AUC of 0.89, 95% CI [0.57, 1.00], compared to a baseline mutation burden AUC of 0.94, 95% CI [0.67, 1.00] for those same samples.

### Predicted Neoantigens are Associated with Mutation Burden and Outcome

In this updated analysis, in contrast to the neoantigen prediction approach employed in (8), NetMHCcons was applied to 8 to 11 amino acid stretches of predicted mutant peptides resulting from both single nucleotide variants (SNV) and indels. All predicted binders less than or equal to 500nM were included, allowing for multiple potential neoepitopes from a single nucleotide variant or indel. Neoepitopes were predicted based on exome data and allele-specific expression was measured in tumors with RNA-Seq data. Tumors had a median of 943 (range 3-6502) predicted neoantigens. Patients who derived clinical benefit had tumors with a higher median predicted neoantigen burden (median 1388, range 209-6502) than those who did not (median 819, range 3-4510) (Mann-Whitney, p=0.01). When considering only pre-treatment tumor samples (n=34), this held true: median predicted neoantigen burden was 1579 (range 209-6502) in benefiters and 582.5 (range 3-2485) in non-benefiters (Mann-Whitney, p=0.002). There was no significant difference in predicted neoantigen burden among post-treatment tumor samples (n=30), with a median predicted neoantigen burden of 940 (range 539-1487) in benefiters and 982 (range 81-4510) in non-benefiters (Mann-Whitney, p=0.49). Using neoantigen burden to predict clinical benefit resulted in an AUC of 0.67, 95% CI [0.54, 0.79] compared to a baseline mutation burden AUC of 0.72, 95% CI [0.58, 0.84]. Considering only pre-treatment samples resulted in an AUC of 0.80, 95% CI [0.64, 0.93] compared to a baseline mutation burden AUC of 0.83, 95% CI [0.69, 0.95] for those samples. There was no difference in the ratio of predicted neoantigens to single nucleotide variants in benefiting tumors versus non-benefiting (median of 2 versus 2.17, respectively). For those pre-treatment tumors with available RNA-Seq data (n=9), the median number of expressed, allele-specific neoantigens was 382.5 (88-725) in benefiters and 48 (1-401) in non-benefiters (Figure 2A, Mann-Whitney, p=0.003), while there was no significant difference in those tumors collected after treatment (Figure 2B, Mann-Whitney, p=0.46). Among pre-treatment tumors (n=9), the AUC for expressed neoantigens was 0.79, 95% CI [0.35, 1.00] compared to a baseline AUC for mutation burden of 0.94, 95% CI [0.67, 1.00]. There was no difference in the percent of expressed predicted neoantigens between benefiting (median 37.4%, range 29.9%-46.3%) and non-benefiting tumors (median 35%, range 14.9%-44.4%).

**Figure 2.**
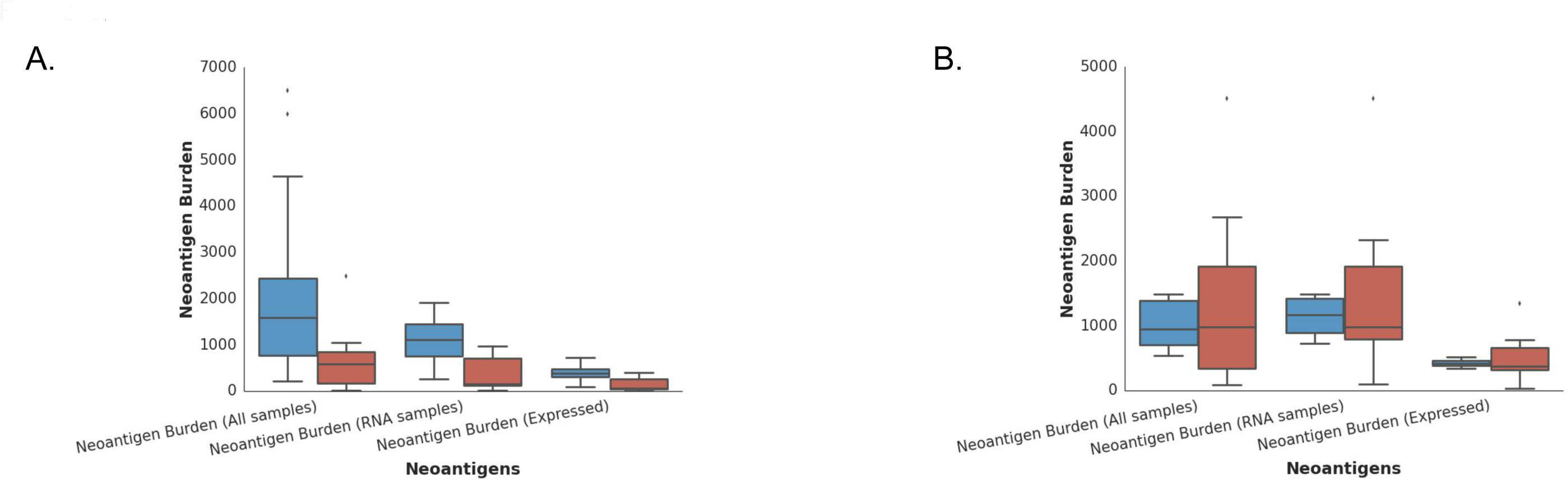
Neoantigen Burden. A. Median and range of neoantigen burden and allele-specific expression of neoantigens in samples collected prior to treatment (for benefit versus no benefit, all (n=34, Mann-Whitney, p=0.002), expressed (n=9, p=0.003)). B. Median and range of neoantigen burden and expressed neoantigens in samples collected after the initiation of treatment (for benefit versus no benefit, all (n=30, Mann-Whitney, p=0.49), expressed (n=15, p=0.46)). In A and B, blue bars represent benefiting tumors; red bars represent non-benefiting tumors. The first pair of bars represent predicted neoantigens for all sequenced tumors; the second pair represent predicted neoantigens in the subset of tumors for which RNA was available; the third represent the expressed neoantigens.

The lack of correlation between benefit and mutation or neoantigen burden in post-treatment samples may suggest immunoediting of specific neoantigens such that the overall neoantigen burden is nearly maintained, but a small number of particularly important neoantigens have been selectively removed.

### Predicted Neoepitope Homology Does Not Outperform Mutation Burden as a Predictor of Response

T cell receptors (TCR) are known to exhibit considerable degeneracy (22), with evidence in infectious diseases that T cells can cross-react to unknown antigens based on homology to antigens to which the host has not previously been exposed (23,24). In cancer, fatalities have been reported that resulted from cross-reactivity of tumor antigen-specific engineered T cell receptors (25). The current RNA-Seq data have been analyzed previously to suggest that an anti-viral interferon-related expression signature is associated with benefit from therapy (12). However, it is unknown whether T cell cross-reactivity plays a role in checkpoint blockade efficacy.

The hypothesis that tumors that respond to checkpoint blockade might harbor recurrent motifs associated with response, either common to responders or homologous to known T cell epitopes remains a question of particular debate. In the initial description of the melanoma sequencing data (8), an algorithm was used to compare 4 amino acid stretches contained within nonamer neoantigens (a “tetrapeptide signature”), irrespective of position or HLA type. We directly evaluated that previous algorithm and separately performed a new comparison.

First, we replicated the tetrapeptide signature approach used by Snyder et al inclusive of the same patients (n=64), variant calls, neoepitope predictions, and IEDB filtering criteria. Unlike Snyder et al, we did not allow held out data to influence the tetrapeptide signature (see Methods). When we categorized tumor samples as featuring or not featuring the tetrapeptide signature, this achieved an AUC of 0.61, 95% CI [0.53, 0.71] compared to an associated mutation burden baseline AUC of 0.76, 95% CI [0.63, 0.89]. Similarly, using counts of signature tetrapeptides rather than a binary representation of signature presence did not outperform mutation burden (Supplementary Information). In addition to directly replicating the approach and data used by Snyder et al, we performed a similar analysis using only pre-treatment, non-discordant samples as well as updated variant calls. The presence of tetrapeptide signature tetrapeptides achieved an AUC of 0.50, 95% CI [0.50, 0.50] compared to an updated mutation burden baseline of 0.85, 95% CI [0.69, 0.98]. Using counts of signature peptides did not outperform mutation burden here, either (Supplementary Information).

In summary, the originally derived tetrapeptide signature and the tetrapeptide signature generated using the revised mutation calling system did not outperform mutation burden as a predictor of clinical benefit, and using the revised system, was not better than random selection.

Next, we created an automated tool, Topeology, for comparing tumor neoepitopes with pathogens from the Immune Epitope Database (IEDB) using sequence alignment of non-anchor residues. This alignment accounts for position, amino acid gaps, and biochemical similarity between amino acids (see Methods). We conducted comparisons of single amino acid substitution-based neoepitopes using this tool, considering only non-discordant, pre-treatment samples (n=33) (Figure 3A-B and Supplementary Figure 2).

**Figure 3.**
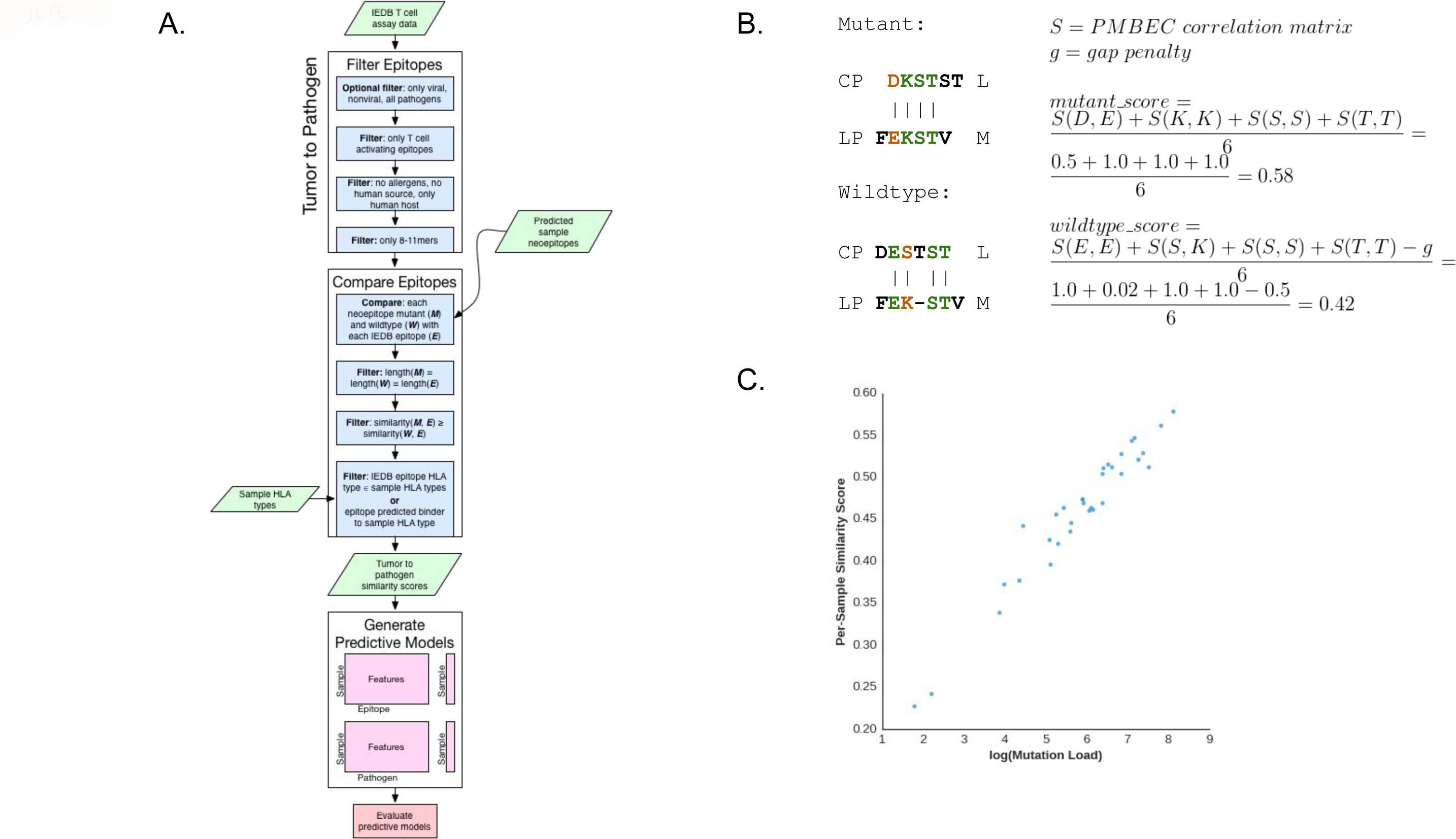
Comparison of Neoepitopes. A. Flow chart illustrating the process for tumor-to-pathogen neoepitope comparison by Topeology. B. An example comparison of the CPDKSTSTL neoepitope (tumor ID 0095) and its wild type peptide to the LPFEKSTVM Influenza A epitope from IEDB. In this case, the neoepitope results in a higher score (0.58) than the wild type peptide (0.42). Only bold amino acids are considered for alignment. Amino acids labeled black do not impact the final Smith-Waterman alignment score for this comparison. Amino acids labeled green are equivalent in both sequences while amino acids labeled orange are not. C. Averaged tumor-to-pathogen similarity scores were highly correlated with the log of mutation burden (Pearson r=0.97, p=6.3e-21).

We explored the extent to which mutation burden may confound these similarity scores. When comparing tumor samples to IEDB pathogens (Supplementary Figure 2), mutation burden was highly correlated with the mean IEDB epitope similarity score for each sample (Figure 3C, Pearson r=0.97, p=6.3e-21). None of these models significantly outperformed the mutation burden baseline AUC of 0.85, 95% CI [0.69, 0.98] (Supplementary Information). We repeated this process using a dataset that includes the neoantigens predicted in a recently published study of 100 tumors from melanoma patients treated with ipilimumab, using the neoantigen predictions and definition of clinical benefit as defined in that study (10). With the caveat that these two neoantigen prediction pipelines were not the same, we found a similar result (Supplementary Information).

Using any of the methods above, the comparison of tumor neoepitopes to pathogens was highly associated with mutation burden but did not outperform mutation burden as a predictor of response.

### Gene Set Enrichment Analysis of Bulk Tumor RNA Expression Illustrates Enrichment for Interferon Signaling and Metabolic Activity in Benefiting Tumors

We sought to determine whether clinical benefit is also associated with an inflamed tumor milieu favorable for anti-tumor immune activation in the setting of CTLA-4 blockade, as previously described (26–29).

When we applied the CIBERSORT method, which was originally developed to deconvolve lymphocyte subsets using microarray data (30), the measure of tumor immune infiltrate correlated with site of origin: resected tumor-containing lymph nodes exhibited a high Pearson correlation coefficient (range: 0.26-0.73) with immune infiltrate as compared to those resected from other primary or metastatic sites (range: -0.04-.32) (per-subtype scores in Supplementary Figure 3). We next applied gene set enrichment analysis (GSEA) using the 50 Hallmark gene sets provided by the MSigDB database (31,32). When we compared all benefiting tumors to non-benefiting tumors, the top 5 most statistically significant gene sets upregulated in benefiting tumors were interferon gamma, interferon alpha, allograft rejection, inflammatory response and complement (Supplementary Figure 4, FDR q-val < 0.005 for each). These data are consistent with previous studies (26). There were no significantly enriched gene sets when all pre-treatment lesions were compared to all post-treatment lesions.

When we examined only pre-treatment tumors (n=9), benefiting tumors (n=4) were characterized by signals of active metabolism, including MTORC1 signaling, glycolysis, and fatty acid metabolism (FDR q-val < 0.05; see Supplementary Information for complete list) relative to non-benefiting (n=5). The UV response (FDR q-val < 0.05) and inflammatory response (FDR q-val < 0.05) gene sets were also significantly enriched.

## Discussion

To date, studies conducted in three tumor types (melanoma (8,10), lung (9) and MSI-high cancers (11)) have illustrated an association between mutation burden and response to checkpoint blockade immunotherapy. Here, we present a reanalysis of previously published data (8) and incorporate new expression data from a subset of patients in that study. Interestingly, predicted Class I candidate neoantigens did not outperform mutation burden as a predictor of response, even when RNA expression was considered. This finding reinforces the importance of other factors to response, including Class II neoepitopes (for which predictive tools remain suboptimal), signaling and cell populations in the tumor microenvironment, and other systemic factors. Furthermore, the ratio of predicted neoantigens to mutations was not significantly different in non-responding tumors, and post-treatment tumors did not exhibit a significant difference in mutation or neoantigen burden between benefiting and non-benefiting tumors. These findings suggest that immunoediting cannot be perceived when evaluating neoantigens in aggregate bioinformatically, but may be occurring at the level of individual neoantigens of particular importance.

In an effort to better evaluate the hypothesis that neoantigens may resemble known pathogens, we created Topeology: a publicly available, biologically relevant tool for peptide comparison to facilitate comparison of T-cell cross-reactivity in any setting (cancer or otherwise). We evaluated neoantigens using both the previously published tetrapeptide signature and Topeology. When either method was applied to two published datasets of melanoma patients treated with CTLA-4 blockade (8,10), we have found that resemblance of neoepitopes to pathogens is associated with but does not outperform mutation burden as a predictor of response to therapy. Therefore, while TCR cross-reactivity may be relevant on an individual T cell level, neither the tetrapeptide signature or neoepitope homology exhibits an indication for use as a biomarker, as both measures are highly associated with, and do not outperform, mutation burden. Topeology may be used to evaluate candidate peptides on an individual basis, for example in the exploration of the hypothesis that tumors generate “danger signals” recognizable to T cells (33).

These data confirm what immunologists have long known (34–36): a myriad of additional factors, ranging from the interferon gamma signaling seen in GSEA analysis (26), to systemic factors (37,38) must be integrated with mutation burden to improve our understanding of tumor response and resistance to checkpoint blockade therapy.

## Acknowledgements

We are grateful for the roles of Vladimir Makarov and Timothy A. Chan in the initial study, to Dr. Makarov in developing the four-caller system, and to Christina Leslie and Jacqueline Buros for their helpful reviews of this manuscript.

Supplementary Information can be found at <u>http://www.hammerlab.org/melanoma-reanalysis</u>

Epitope comparison tool can be found at <u>https://github.com/hammerlab/topeology</u>

## References

1. Hodi FS, O’Day SJ, McDermott DF, Weber RW, Sosman JA, Haanen JB, et al. Improved survival with ipilimumab in patients with metastatic melanoma. N Engl J Med. 2010;363:711–23.

2. Powles T, Vogelzang NJ, Fine GD, Eder JP, Braiteh FS, Loriot Y, et al. Inhibition of PD-L1 by MPDL3280A and clinical activity in pts with metastatic urothelial bladder cancer (UBC). J Clin Oncol. 2014;32:5s.

3. Postow MA, Chesney J, Pavlick AC, Robert C, Grossmann K, McDermott D, et al. Nivolumab and ipilimumab versus ipilimumab in untreated melanoma. N Engl J Med. 2015;372:2006–17.

4. Disis M, Patel MR, Pant S. Avelumab (MSB0010718C), an Anti-PD-L1 Antibody, in Patients with Previously Treated, Recurrent or Refractory Ovarian Cancer: a Phase Ib, Open-label Expansion Trial. J Clin Oncol, Abstract 5509. 2015;

5. Topalian SL, Hodi FS, Brahmer JR, Gettinger SN, Smith DC, McDermott DF, et al. Safety, activity, and immune correlates of anti-PD-1 antibody in cancer. N Engl J Med 2012;366:2443–54.

6. Garon EB, Rizvi NA, Hui R, Leighl N, Balmanoukian AS, Eder JP, et al. Pembrolizumab for the treatment of non-small-cell lung cancer. N Engl J Med. 2015;372:2018–28.

7. Herbst RS, Soria JC, Kowanetz M, Fine GD, Hamid O, Gordon MS, et al. Predictive correlates of response to the anti-PD-L1 antibody MPDL3280A in cancer patients. Nature. 2014;515:563–7.

8. Snyder A, Makarov V, Merghoub T, Yuan J, Zaretsky JM, Desrichard A, et al. Genetic basis for clinical response to CTLA-4 blockade in melanoma. N Engl J Med. 2014;371:2189–99.

9. Rizvi, N., Hellmann, M, Snyder, Et al A. Mutational Landscape Determines Sensitivity to Programmed Cell Death-1 Blockade in Non-Small Cell Lung Cancer. Science. 2015;

10. Van Allen EM, Miao D, Schilling B, Shukla SA, Blank C, Zimmer L, et al. Genomic correlates of response to CTLA-4 blockade in metastatic melanoma. Science. 2015;350:207–11.

11. Le DT, Uram JN, Wang H, Bartlett BR, Kemberling H, Eyring AD, et al. PD-1 Blockade in Tumors with Mismatch-Repair Deficiency. N Engl J Med. 2015;372:2509–20.

12. Chiappinelli KB, Strissel PL, Desrichard A, Li H, Henke C, Akman B, et al. Inhibiting DNA Methylation Causes an Interferon Response in Cancer via dsRNA Including Endogenous Retroviruses. Cell. 2015;162:974–86.

13. Vita R, Zarebski L, Greenbaum JA, Emami H, Hoof I, Salimi N, et al. The immune epitope database 2.0. Nucleic Acids Res. 2010;38:D854–62.

14. Smith TF, Waterman MS. Identification of common molecular subsequences. J Mol Biol. 1981;147:195–7.

15. Kim Y, Sidney J, Pinilla C, Sette A, Peters B. Derivation of an amino acid similarity matrix for peptide: MHC binding and its application as a Bayesian prior. BMC Bioinformatics. 2009;10:394.

16. Berrar D, Flach P. Caveats and pitfalls of ROC analysis in clinical microarray research (and how to avoid them). Brief Bioinform. 2012;13:83–97.

17. Hacohen N, Fritsch EF, Carter TA, Lander ES, Wu CJ. Getting personal with neoantigen-based therapeutic cancer vaccines. Cancer Immunol Res. 2013;1:11–5.

18. Gough CA, Homma K, Yamaguchi-Kabata Y, Shimada MK, Chakraborty R, Fujii Y, et al. Prediction of protein-destabilizing polymorphisms by manual curation with protein structure. PLoS One. 2012;7:e50445.

19. Brash DE. UV signature mutations. Photochem Photobiol. 2015;91:15–26.

20. Cancer Genome Atlas, Network. Genomic Classification of Cutaneous Melanoma. Cell. 2015;161:1681–96.

21. Hugo W, Zaretsky JM, Sun L, Song C, Moreno BH, Hu-Lieskovan S, et al. Genomic and Transcriptomic Features of Response to Anti-PD-1 Therapy in Metastatic Melanoma. Cell. 2016;165:35–44.

22. Adams JJ, Narayanan S, Birnbaum ME, Sidhu SS, Blevins SJ, Gee MH, et al. Structural interplay between germline interactions and adaptive recognition determines the bandwidth of TCR-peptide-MHC cross-reactivity. Nat Immunol [Internet]. 2015; Available from: http://dx.doi.org/10.1038/ni.3310

23. Su LF, Kidd BA, Han A, Kotzin JJ, Davis MM. Virus-specific CD4(+) memory-phenotype T cells are abundant in unexposed adults. Immunity. 2013;38:373–83.

24. Birnbaum ME, Mendoza JL, Sethi DK, Dong S, Glanville J, Dobbins J, et al. Deconstructing the Peptide-MHC Specificity of T Cell Recognition. Cell. 157:1073–87.

25. Cameron BJ, Gerry AB, Dukes J, Harper JV, Kannan V, Bianchi FC, et al. Identification of a Titin-derived HLA-A1-presented peptide as a cross-reactive target for engineered MAGE A3-directed T cells. Sci Transl Med. 2013;5:197ra103.

26. Ji RR, Chasalow SD, Wang L, Hamid O, Schmidt H, Cogswell J, et al. An immune-active tumor microenvironment favors clinical response to ipilimumab. Cancer Immunol Immunother. 2012;61:1019–31.

27. Hamid O, Schmidt H, Nissan A, Ridolfi L, Aamdal S, Hansson J, et al. A prospective phase II trial exploring the association between tumor microenvironment biomarkers and clinical activity of ipilimumab in advanced melanoma. J Transl Med. 2011;9:204.

28. Gajewski TF, Louahed J, Brichard VG. Gene signature in melanoma associated with clinical activity: a potential clue to unlock cancer immunotherapy. Cancer J. 2010;16:399–403.

29. Tumeh PC, Harview CL, Yearley JH, Shintaku IP, Taylor EJM, Robert L, et al. PD-1 blockade induces responses by inhibiting adaptive immune resistance. Nature. 2014;515:568–71.

30. Newman AM, Liu CL, Green MR, Gentles AJ, Feng W, Xu Y, et al. Robust enumeration of cell subsets from tissue expression profiles. Nat Methods. 2015;12:453–7.

31. Subramanian A, Tamayo P, Mootha VK, Mukherjee S, Ebert BL, Gillette MA, et al. Gene set enrichment analysis: a knowledge-based approach for interpreting genome-wide expression profiles. Proc Natl Acad Sci U S A. 2005;102:15545–50.

32. Liberzon A, Subramanian A, Pinchback R, Thorvaldsdottir H, Tamayo P, Mesirov JP. Molecular signatures database (MSigDB) 3.0. Bioinformatics. 2011;27:1739–40.

33. Schumacher T. T Cell Recognition of Human Cancer. 2016.

34. Diamond MS, Kinder M, Matsushita H, Mashayekhi M, Dunn GP, Archambault JM, et al. Type I interferon is selectively required by dendritic cells for immune rejection of tumors. J Exp Med. 2011;208:1989–2003.

35. Kaplan DH, Shankaran V, Dighe AS, Stockert E, Aguet M, Old LJ, et al. Demonstration of an interferon gamma-dependent tumor surveillance system in immunocompetent mice. Proc Natl Acad Sci U S A. 1998;95:7556–61.

36. Nastala CL, Edington HD, McKinney TG, Tahara H, Nalesnik MA, Brunda MJ, et al. Recombinant IL-12 administration induces tumor regression in association with IFN-gamma production. J Immunol. 1994;153:1697–706.

37. Vétizou M, Pitt JM, Daillère R, Lepage P, Waldschmitt N, Flament C, et al. Anticancer immunotherapy by CTLA-4 blockade relies on the gut microbiota. Science. 2015;350:1079–84.

38. Sivan A, Corrales L, Hubert N, Williams JB, Aquino-Michaels K, Earley ZM, et al. Commensal Bifidobacterium promotes antitumor immunity and facilitates anti-PD-L1 efficacy. Science [Internet]. 2015; Available from: http://dx.doi.org/10.1126/science.aac4255

